# Organism-dependent ecotoxicological effects of transition metal-containing layered double hydroxides

**DOI:** 10.64898/2026.11.05.764568

**Authors:** André Lopes Ferreira, Talita Cristina Mena Segato, Camila Carolina de Carvalho, Giovanna Camargo do Carmo, Marcelo A. Mori, Halley Caixeta Oliveira, Rafael Vasconcelos Ribeiro, Marcelo Bispo de Jesus

## Abstract

Due to low environmental stability of certain molecule classes, such as phytohormones and RNA interference, layered double hydroxides (LDHs) have been investigated as nanocarriers capable of protecting and transporting molecules of interest in natural systems. Herein, the ecotoxicological effects of LDH nanoparticles composed by different transition metals (Al-Ni, Al-Co, and Al-Cu) were evaluated in maize (*Zea mays*), tomato (*Lycopersicum esculentum*), and the nematode *Caenorhabditis elegans*, two important crops and one soil organism found in agricultural systems. Plants were exposed to the nanoparticles through foliar spraying (5, 50, and 500 μg mL ¹), and physiological parameters were evaluated. In *C. elegans*, lethality, locomotion, larval development, and body length were analyzed. The nanoparticles exhibited high colloidal stability and characteristic lamellar morphology. The results demonstrated low phytotoxicity for all formulations, without significant changes in photochemical activity, effective quantum yield of photosystem II (ΦPSII), stomatal conductance (gs), and chlorophyll index (Chl a+b; ICF) in plants. In contrast, *C. elegans* exhibited greater sensitivity, showing reduced motility, developmental delay, and decreased body length, although mortality remained below 12% under all evaluated conditions. Toxicity was dependent on the metallic composition, following the order Al-Cu > Al-Ni > Al-Co. Collectively, the results indicate that LDHs exhibit high compatibility with those two crops but may cause sublethal effects in soil organisms. These findings highlight the importance of multispecies ecotoxicological assessments for the safe development of nanocarriers intended for agricultural applications.

**Environmental significance:** Layered double hydroxides (LDHs) have emerged as promising nanocarriers for agricultural applications, particularly for the delivery of environmentally sensitive molecules such as double-stranded RNA (dsRNA). However, their environmental persistence and potential interactions with non-target organisms remain significant factors to consider for their implementation. This study evaluates the ecotoxicological effects of LDH nanoparticles containing various transition metals (Al-Ni, Al-Co, and Al-Cu) on maize, tomato, and the soil nematode *Caenorhabditis elegans*. The results demonstrate high compatibility with both plant species, with no significant impairment to photosynthetic performance or plant growth; conversely, they reveal composition-dependent sublethal effects on the locomotion, development, and body length of the nematodes. The observed toxicity followed the order Al-Cu > Al-Ni > Al-Co, highlighting metallic composition as a key determinant of environmental impact. These findings provide relevant insights into the environmental behavior and biological compatibility of LDH-based nanocarriers, underscoring the importance of multi-species ecotoxicological assessments for the development of safer, nanotechnology-based crop protection strategies.

## 1. Introduction

Ensuring food security remains one of the major global challenges of the twenty-first century. It is estimated that the global population will reach approximately 9.7 billion people by the middle of the century, requiring a considerable expansion of agricultural production to ensure food security^1–3^. Since the World Food Summit in 1996, different policies and programs have been implemented with the aim of increasing agricultural productivity and ensuring access to sufficient and nutritious food for the population^4^. Reinforcing this commitment, the United Nations included the eradication of hunger (SDG 2 – Zero Hunger) among the central goals of the Sustainable Development Goals in 2015. Unfortunately, agriculture still relies on the use of pesticides, playing a fundamental role in the management of organisms capable of compromising agricultural production. The continuous evolution of pesticide-resistant populations, combined with the regulatory withdrawal of active ingredients, stricter requirements regarding chemical residues in agricultural products, and the need to preserve beneficial organisms essential to agroecosystems, has driven the search for innovative and more sustainable crop protection strategies^5^. Consequently, achieving global food security will require crop protection technologies capable of maintaining agricultural productivity while minimizing ecological and human health risks.

In this context, recent advances in nanotechnology and molecular biology have driven a significant transformation in crop protection strategies. In contrast to conventional pesticides—often associated with environmental contamination, biodiversity loss, and the selection of resistant populations nanomaterial-based approaches offer new possibilities for developing control systems that are more specific, efficient, and sustainable. Among these materials, layered double hydroxides (LDHs) are materials composed of hydroxylated layers containing divalent and trivalent metal cations. The partial substitution of M² cations by M³ cations within the layers generates a residual positive charge in the layered structure^6^. To balance this charge and maintain the material’s electrical and structural stability, anionic species along with water molecules are incorporated into the interlayer space. The interaction between the cationic layers and the species within the interlayer region helps maintain overall neutrality and structural organization. LDHs have emerged as some of the most extensively studied nanomaterials for a wide range of applications—including adsorbents, release agents, water electrolysis, wastewater treatment, oil transesterification, agrochemical carriers, and nanocarriers for genes, nanofertilizers, and nanopesticides, due to their desirable physicochemical properties, such as excellent chemical stability, basicity, pH-dependent solubility, and both biological and environmental compatibility^6,7^. Despite these advantages, the environmental fate of LDHs following agricultural application remains poorly understood. Following foliar spraying, a fraction of the applied material may deposit on non-target plants or reach the soil, resulting in the inevitable exposure of non-target terrestrial organisms. Unlike certain molecules such as dsRNA that are highly specific and rapidly degraded under environmental conditions, LDHs consist of more persistent inorganic matrices, the interactions of which with biological systems are strongly influenced by their chemical composition. Consequently, LDHs can remain available for longer periods and interact with a wider range of organisms.

Although LDHs are widely recognized as promising vehicles for agricultural applications, their environmental safety profile remains insufficiently understood. Most available studies have focused on biomedical applications, whereas investigations addressing agricultural exposure scenarios are still scarce. This represents an important knowledge gap, particularly because variations in the divalent and trivalent metal composition of LDHs can substantially alter their physicochemical properties and biological responses. Furthermore, non-target organisms possess distinct physiological traits, biological barriers, and exposure routes, which may result in different susceptibility patterns to the same nanomaterial. Therefore, understanding how different LDH formulations affect non-target organisms is a critical step toward the environmental risk assessment and safe implementation of LDH-based delivery systems in agriculture. Supporting this concern, our previous studies demonstrated that the Al-Cu formulation induced more pronounced sublethal effects than other compositions, likely due to the higher redox activity and biological reactivity associated with Cu² ions^8^.

Therefore, considering that LDHs are expected to persist in the environment longer than the RNA molecules they deliver, this study investigated the ecotoxicological effects of LDH formulations containing different transition metals (Al-Ni, Al-Co, and Al-Cu) on representative non-target terrestrial organisms. Maize (*Zea mays*) and tomato (*Lycopersicum esculentum*) were selected as model crop plants potentially exposed through spray drift and environmental dispersion, whereas *Caenorhabditis elegans* were used as a representative soil nematode to evaluate soil exposure scenarios. We further hypothesized that biological responses would depend on both the metallic composition of the LDH and the exposed organism, with Cu-containing formulations inducing the most pronounced effects^9–11^. To test these hypotheses, LDHs were physicochemically characterized and their effects on plant physiology, as well as nematode lethality, locomotion, development, and body length, were systematically evaluated.

## 2. Materials and Methods

### 2.1 LDH preparation and concentrations

For LDH synthesis, saline solutions (10 mL) were prepared according to a previously described method^12^ using AlCl ·6H O (95%), NiCl ·6H O (99%), CoCl (98%), or CuCl ·6H O (97%). Divalent metal cations (Ni², Co², or Cu²) and Al³ were used at concentrations of 0.70 and 0.30 mol/L, respectively (M² /Al³ ratio of 2.3:1). The mixed solution was slowly added to 40 mL of deaerated NaOH (0.45 mol/L) under nitrogen atmosphere and vigorous stirring for 10 min. The resulting precipitate was collected by centrifugation (6,500 rpm), washed twice with deionized water, and redispersed in 80 mL of deionized water using a vortex. The suspension was subsequently hydrothermally treated at 100 °C for 16 h. Final LDH concentrations were determined gravimetrically by drying 5 mL aliquots overnight at 60 °C, with all exposure concentrations expressed as dry LDH mass per mL of suspension.

### 2.2 Dynamic Light Scattering (DLS) and Stability Assessment

Dynamic light scattering (DLS) was used to determine the hydrodynamic diameter, polydispersity index (PDI), and zeta potential of the nanoparticles using a ZetaSizer Nano ZS90 (Malvern Instruments, Worcestershire, UK). Measurements were performed at 25 °C in disposable polystyrene cuvettes with a 10 mm optical path length and a scattering angle of 90°. Particle size distributions were reported as intensity-weighted distributions. Zeta potential measurements were carried out using folded capillary cells. All LDH suspensions were prepared in ultrapure water at a concentration of 0.1 mg mL ¹, and analyses were performed in triplicate using independent samples. To evaluate pH-dependent stability, the pH of the nanoparticle suspensions was adjusted between 2 and 10 using 0.1 mol L ¹ HCl or NaOH solutions. After 16 h of incubation, changes in hydrodynamic diameter were determined by DLS under the conditions described above.

Thermal stability was assessed using a Litesizer 501 (Anton Paar, Graz, Austria). Nanoparticle suspensions were analyzed in quartz cuvettes (12.5 × 12.5 × 45 mm) over a temperature range of 20-70 °C with temperature increments of 5 °C and an equilibration time of 30 s prior to each measurement. Changes in hydrodynamic diameter were monitored as a function of temperature.

### 2.3 Morphological Characterization

Layered double hydroxide (LDH) nanoparticles were characterized by transmission electron microscopy (TEM) to evaluate their morphology, size, and structural organization. For the analysis, the samples (0.05 mg mL^-1^) were deposited onto copper grids coated with a formvar film and incubated with a 1% (w/v) uranyl acetate solution to provide contrast. Images were acquired using a Tecnai G2 Spirit BioTWIN transmission electron microscope operated at an accelerating voltage of 60 kV.

### 2.4 *Caenorhabditis elegans* and toxicological assays

Wild-type *Caenorhabditis elegans* nematodes were maintained under standard laboratory conditions on plates containing Nematode Growth Medium (NGM) seeded with *Escherichia coli* OP50. Nematode synchronization was performed using the egg-laying technique. Briefly, thirty gravid worms were transferred to each of NGM plates and maintained for 3 h and 30 min to allow egg laying. After this period, the adults were removed from the plates.

For the survival and larval lethality assays, synchronized L4-stage nematodes were washed with 2 mL of M9 buffer and transferred to 15-mL Falcon tubes. Samples were centrifuged at 812 x g for 2 min, the supernatant was discarded, and the pellets were resuspended in 5 mL of M9 buffer. This washing procedure was repeated three times. Subsequently, the pellet was resuspended in 1 mL of M9, and worm density was estimated by counting 10-µL aliquots under a stereomicroscope. Approximately 30 worms per well were distributed into 48-well plates containing a final volume of 1 mL per well, including 100 µL of worm suspension and the compounds diluted in M9 at final concentrations of 5, 50, and 500 µg mL ¹. H O (hydrogen peroxide, 30-volume solution) was used as the positive control, while the negative control consisted only of M9 buffer. All treatments were performed in three biological replicates. After 24 h of exposure, live and dead nematodes were counted. Worms were considered dead when no movement was observed after gentle stimulation of the head or tail using platinum wire picker. The protocol was adapted from Maurer et al. ^13^.

For the body length and locomotion assays, synchronized L4-stage nematodes were transferred to NGM plates containing the nanocarriers at concentrations of 5, 50, and 500 µg mL ¹ and remained exposed for 72 h. For body length analysis, ten worms per treatment were transferred to glass slides containing 10 µL of M9 buffer. Images were acquired using a stereomicroscope and analyzed with ImageJ/FIJI software to determine body length, following previously described methodologies^14,15^. For the locomotion assay, ten worms per treatment were individually evaluated on slides containing 10 µL of M9. Locomotor activity was quantified by counting body bends, defined as movements of the head toward the tail, during a 30-s interval, and the obtained values were multiplied by two to standardize the results as body bends per minute^14,15^.

For the larval development assay, synchronized eggs obtained using the egg-laying method were directly exposed to the tested nanocarriers incorporated into NGM at concentrations of 5, 50, and 500 µg mL ¹, allowing exposure from the embryonic stage onward. After 60 h of exposure (since embryogenesis), the number of nematodes reaching adulthood was monitored over the subsequent 12 h. Adult development was determined by the appearance of eggs in the gonad, as previously described by Corsi et al. (2018) ^16^. The percentage of animals reaching adulthood was calculated for each treatment and compared with the control groups. Results were expressed as mean ± standard error of the mean (SEM) and statistically analyzed using Scheffé’s test.

### 2.5 Crop species, foliar spraying, physiological and growth evaluations

Tomato plants (*Lycopersicum esculentum*, IKRAM F1 Hybrid) were cultivated in plastic pots (3 L) filled with a commercial substrate containing rice husk, *Sphagnum* spp., and perlite in a 7:2:1 ratio,pH 5.5; Electrical Conductivity 0.7 mS/cm; Max. Moisture 70% w/w. (Carolina Soil of Brazil, Vera Cruz RS, Brasil). The plants were grown in a greenhouse, where air temperature and relative humidity were continuously monitored. The average minimum and maximum air temperatures were 20 °C and 38 °C, respectively, while air relative humidity ranged from 52% to 89%. The photoperiod averaged approximately 12 h, and the photosynthetic photon flux density (PPFD) reached approximately 1,100 µmol m ² s ¹ near midday under clear-sky conditions.The plants were irrigated daily throughout the experimental period. Sixteen days after sowing, the plants were fertilized with 100 mL of a modified Sarruge (1975) nutrient solution containing: N (16%), P O (8%), K O (12%), MgO (2%), S (5%), Fe (0.4%), B (0.02%), Zn (0.02%), Cu (0.05%), Mn (0.06%), and Mo (0.015%), as done previously^17^.

The experiment was conducted in Londrina, Paraná, Brazil. Hybrid maize seeds *(Zea mays* L. cv. Balu 787) were directly sown in conical high-density polypropylene tubes (290 mL) filled with the commercial substrate Carolina Soil® (formulation 75H, Class LXXVI; Carolina Soil of Brazil, Vera Cruz, RS, Brazil), composed of *Sphagnum* peat, expanded vermiculite, dolomitic limestone, agricultural gypsum, and NPK fertilizer (pH 5.5, Electrical Conductivity 0.7 mS/cm) and grown in a greenhouse covered with a shade screen with 20% light retention (i.e., ∼80% light transmittance), under natural temperature, relative humidity, and photoperiod conditions (day length ≈ 14 h during the experimental period) throughout the experimental period. Plants were irrigated daily to maintain adequate substrate water availability. No post-emergence fertilization was applied, as the substrate already contained NPK fertilizer.

In a completely randomized design, three formulations of metallic nanoparticles (Al-Ni, Al-Co, and Al-Cu) were sprayed at three concentrations (5, 50, and 500 µg mL^-1^). Nanoparticle stock suspensions were diluted in distilled water to obtain the concentration used in the study. In maize, foliar spraying was done 11 days after sowing, when plants reached the V2 phenological stage. Before spraying, homogenous plants were selected based on their size and developmental stage to reduce experimental variability. The plants were sprayed a single time with a nanoparticle suspension diluted in water, using a total volume of 1 mL per plant. Control plants received the same volume of distilled water. The suspensions were sprayed at 9:00 a.m. onto the adaxial and abaxial leaf surfaces to ensure uniform coverage.

Measurements of leaf gas exchange in maize plants were performed on the first and sixth days after foliar spraying, using a portable infrared gas analyzer (LI-6400XT model; LI-COR Biosciences, Lincoln NE, USA). Net photosynthetic rate (*A*, μmol m ² s ¹), transpiration (*E*, mmol m^-2^ s^-1^), intercellular CO_2_ concentration (*C*_i_), and stomatal conductance (*g*_s_, mol H_2_O m^-2^ s^-1^) were measured in fully expanded leaves using a 6-cm² chamber under saturating photosynthetically active radiation (1,500 μmol m ² s ¹) and an air flow rate of 500 μmol s ¹. Measurements were performed on sunny days starting at 8:00 and 11:00 a.m. During the first measurement (first day after spraying), air temperature ranged from 29.9 to 39.1 °C, relative humidity ranged from 27.3% to 41.5%, and reference CO_2_ concentration ranged from 391.7 to 407.8 μmol mol^-1^. On the second measurement (sixth day after spraying), air temperature ranged from 30.7 to 37.9 °C, relative humidity ranged from 32.6% to 47.3%, and reference CO_2_ concentration ranged from 393.4 to 408.9 μmol mol^-1^, as recorded by the LI-6400XT. The potential quantum efficiency of photosystem II (F_v_/F_m_= (F_m_-F_0_)/F_m_) was determined in intact leaves using an OS1p fluorometer (Opti-Sciences, USA), the minimum fluorescence (F_0_) was measured using a weak modulated light, and maximum fluorescence (F_m_) was determined after 0.8-s saturating light pulse of 8250 μmol m ² s ¹. Readings were taken early in the morning, between 6:00 and 8:00 a.m., 15 min after dark adaptation of the leaves using FL-DC clips (Opti-Sciences). The experiment ended 10 days before the formulation applications (21 days after sowing). At the end of the experiments, plant growth was evaluated by measuring stem and root lengths using a ruler. Stem, leaf, and root tissues were then collected separately, placed in paper bags, and dried in an oven at 60 °C for 72 h to determine their respective dry masses.

In tomato plants, simultaneous measurements of the effective quantum efficiency of PSII (Φ_PSII_) and stomatal conductance (*g*_s_, mol H_2_O m^-2^ s^-1^) were taken in fully expanded and mature leaves under natural conditions (light, air temperature and relative humidity) with a porometer/fluorometer model LI-600 (LI-COR Biosciences, Lincoln NE, USA). Measurements were done between 09:00 and 11:00 a.m. and data were recorded after temporal stability, following the manufacturer recommendations and using the Auto mode. After measuring photochemical activity and stomatal aperture, we used a chlorophyll meter (ClorofiLOG CFL1030, Falker, Porto Alegre RS, Brazil) to evaluate the content of chlorophylls *a* and *b* and then *a+b*, as shown herein as chlorophyll index (CCI).

### 2.6 Statistical analysis

All data were analyzed using GraphPad Prism software (version 8.0, GraphPad Software, San Diego CA, USA). Results are presented as mean ± standard error of the mean (SEM). Differences among treatments were evaluated by one-way analysis of variance (ANOVA) followed by Scheffé’s multiple comparison test. Mortality data were analyzed using nonlinear regression with a four-parameter logistic dose–response model, plotting the logarithm of nanoparticle concentrations to generate sigmoidal curves. When appropriate, comparisons were made between treated groups and the control group, as well as among concentrations within the same compound. Statistical significance was considered at p < 0.05.

## 3. Results

### 3.1 Physicochemical Stability of LDH Nanoparticles Under Agriculturally Relevant Conditions

We monitored the physicochemical stability of LDH formulations to determine their stability under storage and environmental fluctuations. Hydrodynamic diameter and zeta potential were periodically assessed over 12 months at room temperature. As shown in Figure 1B-C, Al-Ni and Al-Co nanoparticles maintained stable size and surface charge throughout the monitoring period, indicating strong long-term aqueous stability. Al-Cu also exhibited good colloidal stability, with only minor variations in physicochemical parameters over time.

**Figure 1.**
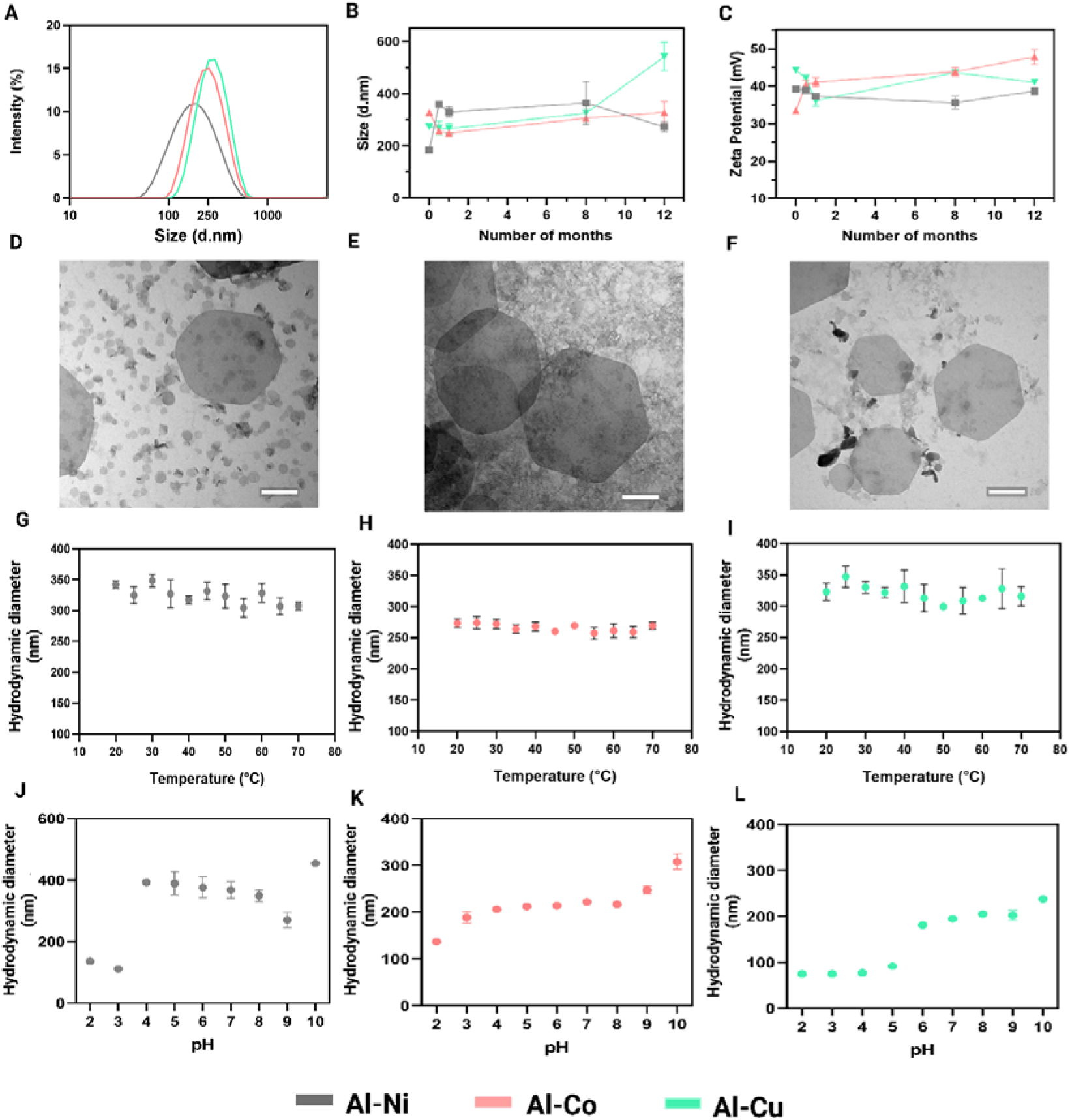
Physicochemical characterization and stability assessment of Al-Ni, Al-Co, and Al-Cu layered double hydroxide (LDH) nanoparticles. (A) Hydrodynamic size distribution determined by dynamic light scattering (DLS). (B) Hydrodynamic diameter and (C) zeta potential measured during 12 months of storage. (D-F) Transmission electron microscopy (TEM) images of Al-Ni, Al-Co, and Al-Cu nanoparticles, respectively. Scale bar = 200 nm. (G-I) Hydrodynamic diameter of Al-Ni, Al-Co, and Al-Cu nanoparticles, respectively, measured at different temperatures (20-70 °C). (J-L) Hydrodynamic diameter of Al-Ni, Al-Co, and Al-Cu nanoparticles, respectively, measured under different pH conditions (pH 2-10). Hydrodynamic diameter and zeta potential were determined by dynamic light scattering (DLS).

Transmission electron microscopy (TEM) micrographs confirmed the formation of layered double hydroxide (LDH) nanoparticles with the characteristic lamellar morphology for all compositions evaluated (Figure 1D-F). Al-Ni nanoparticles (Figure 1 D) exhibited a predominantly hexagonal morphology with a relatively heterogeneous size distribution, including both isolated particles and small aggregates. Al-Co formulations (Figure 1 E) displayed well-defined hexagonal platelets with larger lateral dimensions while preserving the typical layered structure of LDHs. Similarly, Al-Cu nanoparticles (Figure 1 F) showed a preserved hexagonal morphology, with individualized particles and clearly defined edges. For all formulations, the TEM images revealed particles with a high lateral dimension-to-thickness ratio, consistent with the layered architecture of LDHs, and no evidence of significant morphological alterations or detectable secondary phases. These results confirm the successful synthesis of LDH nanoparticles and the preservation of their lamellar structure regardless of the divalent metal cation employed in the synthesis.

To assess resilience to temperature variation, we further evaluated hydrodynamic diameter under thermal exposure. DLS measurements revealed no significant changes across the tested temperature range for Al-Ni, Al-Co, and Al-Cu formulations, demonstrating that thermal fluctuations did not induce particle aggregation or structural degradation.

Overall, these results demonstrate that Al-Ni, Al-Co, and Al-Cu LDH nanoparticles exhibit high colloidal and thermal stability, supporting their suitability for agricultural and biotechnological applications that require stability under prolonged storage and variable environmental conditions

The thermal degradation analysis of Layered Double Hydroxide (LDH) nanoparticles composed of Al-Ni, Al-Co, and Al-Cu, monitored by Dynamic Light Scattering (DLS), showed no significant changes in hydrodynamic size as a function of temperature for any of the compositions (Figure 1 G-I). These findings indicate that all formulations remained structurally stable throughout the thermal assay. The evaluation of pH effects on colloidal behavior revealed that the hydrodynamic size of the nanoparticles varies in a pH-dependent manner (Figure 1 J-L). For the Al-Ni LDH, a marked increase in hydrodynamic diameter was observed between pH 4 and 7, whereas at pH 2-3 the particles remained smaller and more dispersed. The Al-Co LDH showed greater stability between pH 3 and 7, with only minor variations in mean particle size. In contrast, the Al-Cu LDH exhibited smaller sizes under acidic conditions and a progressive increase above pH 6.

### 3.2 Physiological responses of non-target plants to LDH exposure

#### 3.2.1 Phytotoxicity of LDH Nanoparticles to Maize Plants

Foliar spraying of LDH nanoparticles (Al-Ni, Al-Co, and Al-Cu) at concentrations of 5, 50, and 500 µg mL ¹ did not significantly affect leaf gas exchange or primary photochemistry up to 6 days after application (Figure 2). The net photosynthesis rate (A) remained relatively stable across treatments and concentrations, ranging approximately from 28 to 40 µmol m ² s ¹. Similarly, stomatal conductance (gs) showed moderate variation among treatments, ranging approximately from 0.21 to 0.48 mol H O m ² s ¹, but without a consistent pattern dependent on treatment or concentration. Overall, no statistically significant differences in A or gs relative to the control were detected throughout the evaluated period (Figure 2). In maize plants, foliar application of LDH nanoparticles (Al-Ni, Al-Co, and Al-Cu) did not promote marked changes in internal CO concentration (Ci) or transpiration rate (E) compared with the control (Figure S1). The broad overlap of standard deviations between treated groups and the control suggests that LDH nanoparticles did not significantly affect leaf gas exchange in maize plants under the evaluated experimental conditions. Accordingly, the maximum quantum efficiency of photosystem II (Fv/Fm) remained stable across all treatments and evaluation times (Figure 2 H-J), varying between 0.72 and 0.76.

**Figure 2.**
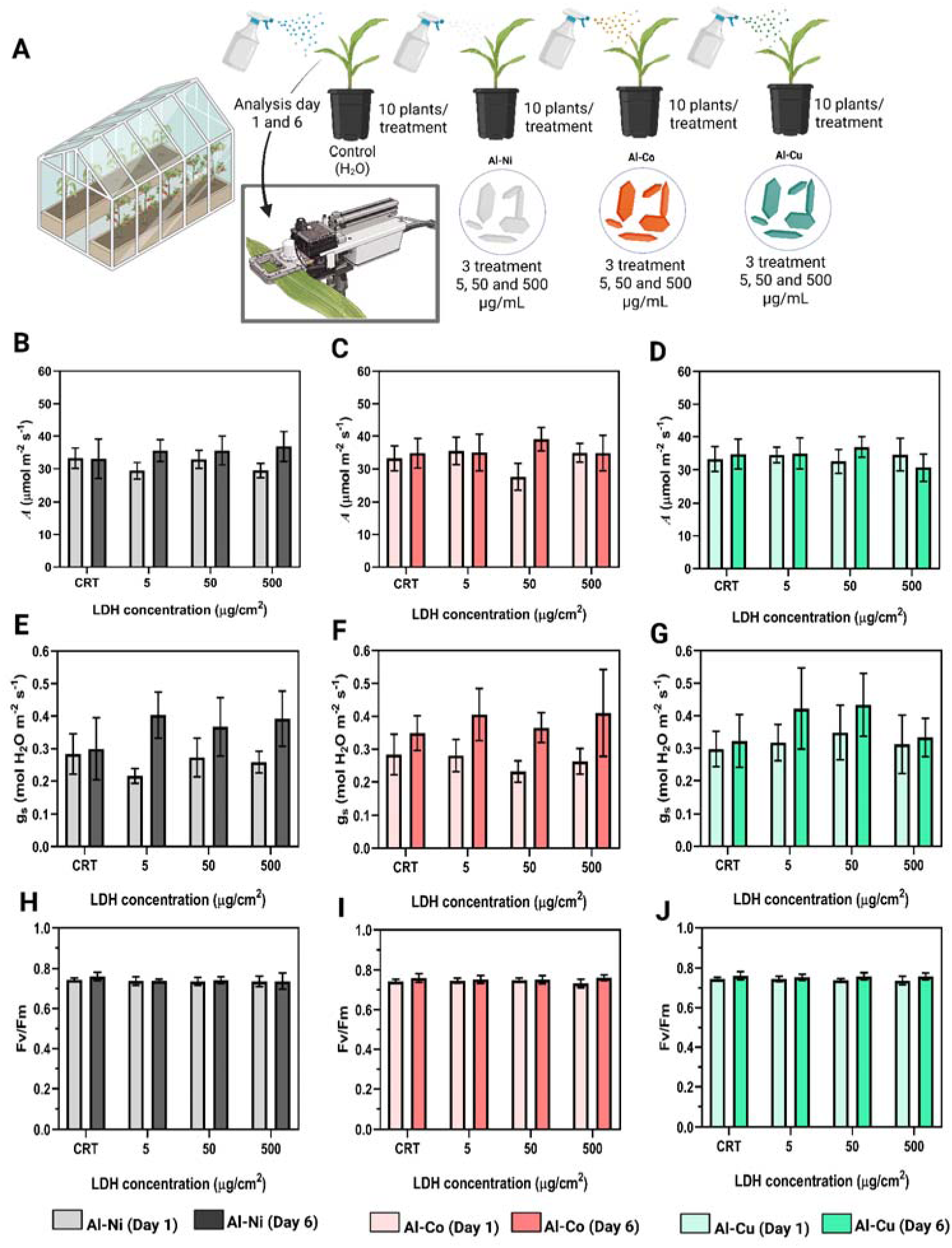
Experimental design and physiological responses of maize plants exposed to layered double hydroxide (LDH) nanoparticles with different metal compositions (Al-Ni, Al-Co, and Al-Cu). (A) Schematic representation of the greenhouse experiment. Maize plants were sprayed with LDH suspensions at concentrations of 5, 50, and 500 μg mL ¹, with water-treated plants serving as controls (CRT). Leaf gas exchange and photochemistry were evaluated 1 and 6 days after spraying plants. (B-D) Net photosynthetic rate (*A*), (E-G) stomatal conductance (*g*_s_), and (H-J) maximum quantum efficiency of photosystem II (F_v_/F_m_) measured in plants treated with Al-Ni, Al-Co, and Al-Cu LDHs, respectively. Data are presented as mean ± standard deviation (n = 7 plants per treatment).

To better assess whether the foliar application of LDH nanoparticles affected plant growth, stem and root lengths were analyzed at the end of the experiments. No statistically significant differences were observed among treatments for either root or stem length parameters, regardless of nanoparticle composition or applied concentration (Figure S2 A–F). Similarly, stem, leaf, and root dry masses remained comparable to those of control plants across all treatments, with no statistically significant changes associated with LDH application (Figure S3 A–I). These results indicate that the foliar application of Al-Ni, Al-Co, and Al-Cu LDH nanoparticles did not interfere with the growth or biomass accumulation of maize plants under the experimental conditions evaluated.

Overall, the physiological parameters evaluated up to 6 days after foliar spraying remained similar to those observed in control plants, regardless of the metallic composition of the nanoparticles or the concentration used. No evidence of impairment in photosynthetic activity and stomatal regulation was observed under the experimental conditions evaluated.

#### 3.2.2 Assessment of the phytotoxicity of LDH nanoparticles in tomato plant

Foliar spraying of LDH nanoparticles (Al-Ni, Al-Co, and Al-Cu) to tomato plants did not promote marked changes in photochemistry, pigments and stomatal conductance evaluated throughout the experimental period (Figure 3). Overall, the observed values remained close to those of the control group, regardless of the metallic composition of the nanoparticles or the applied concentration. The effective quantum yield of photosystem II (Φ_PSII_) showed only minor variations among treatments, remaining close to the values observed in control plants and ranging between 0.60 and 0.77 at the 1^st^ day and between 0.57 and 0.62 at the 6^th^ day (Figure 3 B-D). No consistent concentration-dependent effect was observed for any of the evaluated formulations. In accordance, the chlorophyll index (given by Chl *a+b*, CCI) remained high across all treatments, varying between 39 and 47 (Figure 3 H-J). For the Al-Ni and Al-Cu formulations, mean values were similar to those of the control at all evaluated concentrations. Chlorophyll indices (Chl *a*, Chl *b*, and Chl *a+b*) also remained relatively stable in tomato plants treated with Al-Ni, Al-Co, and Al-Cu LDH nanoparticles, with values comparable to those of the control across all evaluated concentrations (Figure S4 A–I). Overall, these results indicate that foliar application of the different LDH formulations did not affect leaf pigment maintenance or photosynthetic capacity during the experimental period, in agreement with the chlorophyll fluorescence and gas exchange results. Together, these results indicate that foliar application of the LDH formulations did not impair photochemical activity in tomato plants during the experimental period. Stomatal conductance (g_s_) also remained relatively stable among treatments (Figure 3 E-G). Plants treated with Al-Ni exhibited values ranging from 0.45 to 0.68 mol H O m ² s ¹, whereas treatments with Al-Co and Al-Cu showed values varying from 0.50 to 0.66 mol H O m ² s ¹. Although small fluctuations were observed between evaluation times, no pattern associated with nanoparticle concentration was identified.

**Figure 3.**
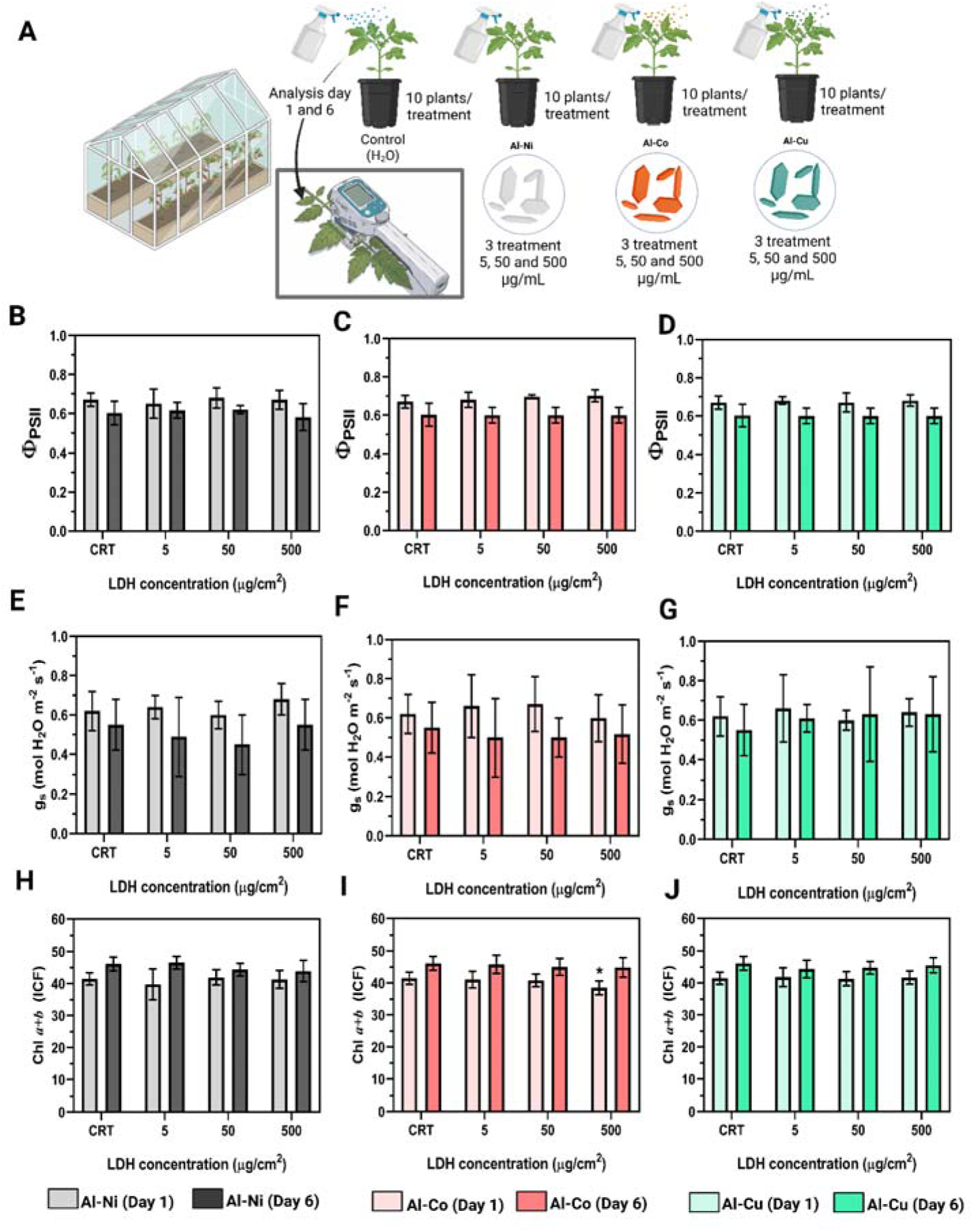
Experimental design and effects of layered double hydroxide (LDH) nanoparticles on photosynthetic performance and chlorophyll content of tomato plants. (A) Schematic representation of the greenhouse experiment. Tomato plants were foliar-sprayed with LDH nanoparticles containing different metal compositions (Al-Ni, Al-Co, and Al-Cu) at concentrations of 5, 50, and 500 μg mL ¹, while water-treated plants served as controls (CRT). Ten plants were used per treatment, and physiological measurements were performed 1 and 6 days after foliar spraying. (B-D) Effective quantum yield of photosystem II (Φ_PSII_), (E-G) stomatal conductance (g_s_), and (H-J) chlorophyll index (Chl a+b; ICF) measured in plants treated with Al-Ni, Al-Co, and Al-Cu LDHs, respectively. Data are presented as mean ± standard deviation (n = 10 plants per treatment). Asterisks indicate statistically significant differences compared with the respective control treatment (*p < 0.05).

Overall, the results demonstrate that foliar application of LDH nanoparticles did not compromise the photochemical efficiency of photosystem II, chlorophyll content and stomatal aperture in tomato plants during the evaluated period.

### 3.3 Effect of LDH Nanoparticles on the Growth and Development of *C. elegans*

Exposure of *C. elegans* to LDH nanoparticles for 24 h resulted in low levels of lethality under all evaluated conditions (Figure 4 B). For the Al-Co compound, mortality rates were 6.3%, 5.8%, and 8.3% at concentrations of 5, 50, and 500 µg mL ¹, respectively. Treatment with Al-Cu resulted in lethality of 6.5%, 5.6%, and 4.0%, whereas the Al-Ni compound induced mortality rates of 1.2%, 3.8%, and 11.1% at concentrations of 5, 50, and 500 µg mL ¹, respectively. In none of the treatments was it possible to estimate the LC within the evaluated concentration range (5-500 µg mL ¹), since mortality remained below 50% under all experimental conditions.

**Figure 4.**
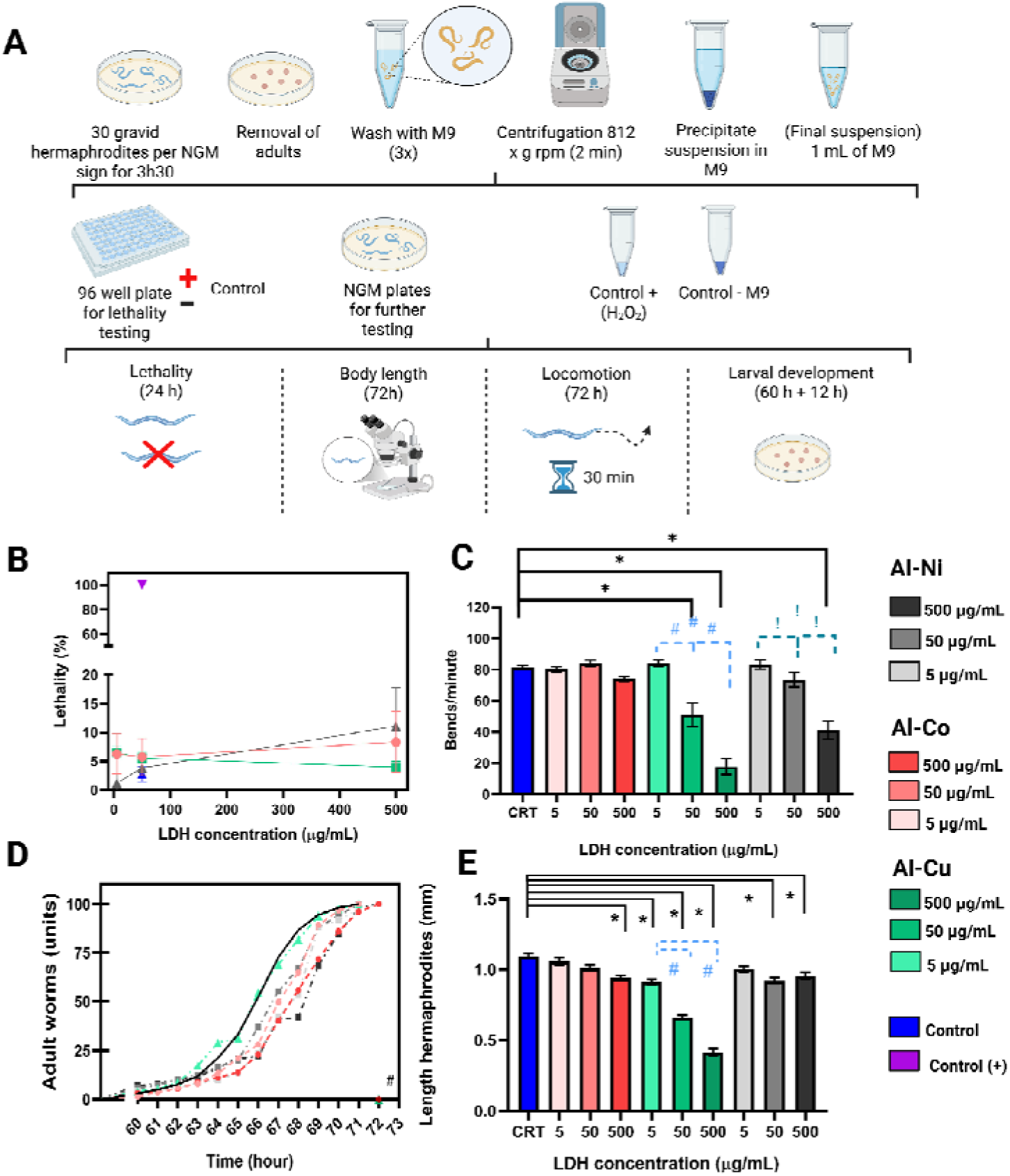
Evaluation of the ecotoxicological effects of LDH nanoparticles on *Caenorhabditis elegans*. (A) Schematic representation of the experimental design used for lethality, locomotion, larval development, and body length assays. (B) Percentage lethality of *C. elegans* exposed to Al-Co, Al-Cu, and Al-Ni formulations at concentrations of 5, 50, and 500 µg mL ¹ after 24 h of exposure, using 30 nematodes per well, with three technical replicates per treatment (n=3). (C) Quantification of locomotion, expressed as body bends per minute, after 72 h of exposure to the treatments, using 10 nematodes per well, with three technical replicates per treatment (n=3). (D) Larval development of *C. elegans* after 60 h of exposure to the different LDH formulations, using 100 nematodes per treatment (n=3). (E) Body length of nematodes after 72 h of exposure to the treatments, using 10 nematodes per well, with three technical replicates per treatment (n=3). Data are presented as the mean ± standard error of the mean (SEM). *p* < 0.05 compared with the control group; # *p* < 0.05 among concentrations within the Al-Cu treatment; ! *p* < 0.05 among concentrations within the Al-Ni treatment.

The effects of the compounds on nematode locomotion after 72 h of exposure. Treatment with Al-Cu promoted a significant reduction in locomotion at concentrations of 50 and 500 µg mL ¹ compared with the control group. For the Al-Ni compound, a significant decrease was observed only at the concentration of 500 µg mL ¹ (Figure 4 C).

The developmental assessment after more than 60 h of exposure to Al-Co at 500 µg mL ¹ and to Al-Cu at concentrations of 50 and 500 µg mL ¹ resulted in the presence of individuals in the dauer stage, evidence of a stress condition. In addition, except for the Al-Cu treatment at 5 µg mL ¹, all exposed groups exhibited developmental delay compared with the control (Figure 4 D).

Changes in nematode body length after 72 h of exposure the Al-Co compound promoted a significant reduction in body length only at the concentration of 500 µg mL ¹. For the Al-Cu treatment, significant reductions were observed at all evaluated concentrations (5, 50, and 500 µg mL ¹). The Al-Ni compound resulted in a significant decrease in body length at concentrations of 50 and 500 µg mL ¹ (Figure 4 E).

## 4. Discussion

The environmental fate of nanoparticle-based delivery systems is a critical aspect for the development of sustainable agricultural technologies. Unlike conventional active ingredients, nanocarriers may persist in the environment after the release of the carried molecule, potentially increasing the exposure of non-target organisms. Furthermore, it is essential to evaluate the potential phytotoxicity of these nanomaterials in the treated plants, since adverse effects on plant physiology or photosynthetic performance could compromise crop health and limit the safe application of nanotechnology-based agricultural technologies. Therefore, understanding the biological effects associated with the nanocarrier itself is essential for a comprehensive environmental risk assessment. In this study, we established an experimental framework to evaluate the ecotoxicological and physiological responses of representative non-target organisms occupying different ecological niches. The soil-dwelling nematode *Caenorhabditis elegans* was selected as a model organism to assess potential impacts on soil biota, while maize (*Zea mays*) and tomato (*Lycopersicum esculentum*) were used as non-target plant models to investigate possible effects on plant physiology. This approach allows the evaluation of whether LDH nanoparticles remain biologically compatible with key terrestrial organisms under conditions relevant to agricultural applications^18,19^.

The metallic composition of LDH nanoparticles emerged as a key determinant of their biological effects, highlighting the importance of metal selection in the design of environmentally safe nanocarriers. Despite showing no phytotoxic effects in the plant models evaluated, assays conducted with *Caenorhabditis elegans* revealed clear differences in toxicity among the formulations. Overall, the results established a toxicity ranking of Al-Cu > Al-Ni > Al-Co, indicating that the substitution of the divalent metal substantially influences nanoparticle–organism interactions. The greater toxicity associated with Al–Cu nanoparticles may be related to the well-documented biological reactivity of copper, which can promote oxidative stress, disrupt cellular homeostasis, and interfere with essential metabolic pathways^20^. In contrast, Al–Co nanoparticles induced comparatively weaker biological responses, suggesting lower bioavailability or reduced interaction with biological targets. These findings indicate that, although the LDH structure remains similar among formulations, variations in metallic composition can alter their environmental behavior and toxicological profile. Therefore, the selection of the divalent metal should be considered a critical parameter in the development of LDH-based delivery systems intended for agricultural applications.

The induction of dauer arrest in *C. elegans* exposed to Al-Co (500 µg mL ¹) and Al-Cu (50 and 500 µg mL ¹) provides a mechanistically informative organism-level endpoint that extends beyond the descriptive sublethal alterations observed in this study. Dauer entry is a genetically regulated developmental decision governed by the integration of insulin/IGF-1 signaling (DAF-16/FoxO), steroid hormone signaling (DAF-12/NHR), and TGF-β pathways, and is induced in response to adverse environmental conditions, including oxidative and other cellular stresses, that favor developmental arrest over continued growth^21^. Notably, the concentration threshold for dauer induction followed the same ranking observed for the other sublethal endpoints, with Al-Cu inducing the response at lower concentrations than Al-Co, suggesting that both outcomes may arise from activation of a common stress-signaling axis^22^. Likewise, CuO nanoparticles have been reported to reduce body length and induce neurodegeneration in this model^23^. Although the present study did not directly assess reactive oxygen species (ROS) production or DAF-16 activation, the induction of dauer represents an integrated organism-level stress response. Taken together, the concordant concentration-dependent effects on body growth, locomotion, developmental progression, and dauer induction suggest that these phenotypes represent successive manifestations of a common physiological stress response.

A key finding of this study is that biological responses to LDH exposure varied among the tested organisms. In maize and tomato plants, foliar application of Al-Ni, Al-Co, and Al-Cu-containing LDH nanoparticles did not significantly affect the evaluated photosynthetic parameters or chlorophyll content under the tested conditions. In contrast, *C. elegans* exhibited measurable sublethal effects within the same concentration range. These differences may reflect distinct exposure pathways, uptake mechanisms, and biological barriers among the tested organisms^24,25^, whereas soil nematodes may experience more direct interactions with nanomaterials through contact and ingestion. From an ecotoxicological perspective, these findings highlight the importance of evaluating endpoints beyond mortality when assessing the environmental safety of nanomaterials. Changes in locomotion, growth, and development may influence individual fitness and potentially affect ecological functions performed by soil organisms, even when acute lethality is low. Collectively, the results indicate that the biological effects of LDH nanoparticles are species– and composition-dependent, reinforcing the need for multispecies approaches in environmental risk assessment.

In theory, nanoparticle exposure may disrupt cellular redox homeostasis, increase ROS accumulation, and interfere with cell cycle progression, potentially causing cytogenotoxic damage to proteins, lipids, and nucleic acids and ultimately affecting plant development and physiological performance^26^. However, in the present study, LDH nanoparticles containing nickel, cobalt, or copper exhibited low phytotoxicity in maize and tomato plants, even at the highest concentration tested (500 µg mL ¹). Physiological parameters remained within the range observed in control plants, with no significant effects on photosynthetic activity or stomatal dynamics.

The absence of significant phytotoxic effects may be related to the physicochemical properties of LDHs and the biological barriers regulating nanoparticle uptake^24^. Nanoparticle bioavailability is influenced by particle size, surface charge, aggregation, and dissolution rate^27^. Alternatively, metallic species associated with nanoparticles may have been compartmentalized or precipitated before inducing detectable changes in chloroplast photochemistry. Further studies addressing nanoparticle uptake, translocation, and fate within plants are therefore needed, particularly for crops such as maize and tomato that produce edible seeds and fruits.

Although the release of metal ions was not directly quantified in this study, the low phytotoxicity observed in corn and tomato plants may be associated with the intrinsic physicochemical properties of LDHs (layered double hydroxides) specifically their lamellar structure, gradual dissolution, and low bioavailability^28^. The high colloidal stability demonstrated by the LDH formulations may further contribute to their biological compatibility by limiting uncontrolled dissolution and the release of high concentrations of free metal ions factors frequently linked to phytotoxic responses. In this context, the confinement of metal species within the LDH structure, combined with the limited uptake of nanoparticles due to plant barriers, may explain the absence of significant phytotoxic effects, rather than this being attributable to the intrinsic biological role of the constituent metals^25^. Furthermore, the physicochemical robustness of these formulations—reflected in their long-term colloidal stability and resistance to temperature fluctuations—supports their potential use as environmentally compatible nanocarriers in agricultural applications that require prolonged storage and exposure to variable environmental conditions.

On the other hand, assays conducted under varying pH conditions demonstrated that the colloidal behavior of the formulations was dependent on the chemical conditions of the medium. Al-Ni and Al-Cu nanoparticles exhibited smaller hydrodynamic diameters under acidic conditions, a behavior frequently associated with the protonation of hydroxyl groups present in the LDH layers and the consequent partial dissolution of the structure. This process may favor the gradual release of constituent metal ions, increasing their bioavailability in biological environments. This hypothesis is particularly relevant for interpreting the results obtained with *C. elegans*. Although mortality remained low under all evaluated conditions, significant sublethal effects on nematode locomotion, development, and body growth were observed. The greater toxicity observed for the Al-Cu formulation may be related to the higher biological reactivity of Cu² ions released following partial nanoparticle dissolution. Copper is widely recognized for its pro-oxidant potential and its ability to induce metabolic alterations, mitochondrial dysfunction, and oxidative stress in different organisms^26^. Therefore, the combination of partial LDH dissolution under specific environmental conditions and the high biological activity of copper may help explain the greater sensitivity observed in organisms exposed to the Al-Cu formulation.

These results reinforce that the physicochemical stability of nanomaterials and their response to different environmental conditions should be considered together with biological factors to achieve a more comprehensive understanding of their ecotoxicological impacts. The metals present in LDH structures play different physiological roles in plants, which may influence their interaction with biological systems. Nickel is recognized as an essential micronutrient, acting as a cofactor for several enzymes related to nitrogen metabolism, cellular detoxification, and maintenance of redox homeostasis^29^. Copper also plays a fundamental role in plant development, participating in processes such as mitochondrial respiration, photosynthesis, cellular signaling, and antioxidant defense^30–33^. In contrast, cobalt is not considered an essential element for most plant species, although it may exert beneficial effects under certain conditions. Its uptake and distribution within plant tissues depend on transport mechanisms shared with other divalent cations and are regulated by specific physiological pathways

Although metal-ion release was not directly quantified, the results suggest that pH-dependent dissolution may contribute to the toxicity differences among the evaluated formulations. Future studies should therefore quantify metal dissolution and assess oxidative stress biomarkers to clarify the mechanisms involved. Several limitations should also be acknowledged. The present experiments focused on short-term responses and did not address chronic or repeated exposure under environmentally relevant conditions. Although Caenorhabditis elegans showed formulation-dependent toxicity, the relative contributions of nanoparticle-specific effects and dissolved metal ions remain unclear. In addition, nanoparticle uptake, translocation, and accumulation in plant tissues, as well as potential effects on soil physicochemical properties and other non-target organisms, were not evaluated. Future research integrating mechanistic endpoints, such as ROS production and molecular stress responses, with broader ecotoxicological assessments will be important for clarifying the environmental fate and safety of LDH-based nanocarriers in agricultural systems.

## 5. Conclusion

Our results demonstrate that the biological responses to Al–Ni, Al–Co, and Al–Cu LDH nanoparticles were dependent on the exposed organism, metallic composition, and concentration. Physicochemical characterization confirmed the successful formation of LDH nanoparticles with the characteristic lamellar morphology and high lateral dimension-to-thickness ratio. The three formulations also exhibited good long-term colloidal stability over 12 months, while their response to temperature variation indicated high thermal stability without significant changes in hydrodynamic size. Although colloidal behavior varied as a function of pH and metallic composition, the observed stability under the tested conditions highlights the physicochemical robustness of these materials. In *Caenorhabditis elegans*, the formulations induced distinct sublethal effects, establishing a toxicity ranking of Al–Cu > Al–Ni > Al–Co. In contrast, maize and tomato plants showed high tolerance to nanoparticle exposure, with no significant physiological impairments observed under the tested conditions.

The differences observed among formulations highlight metallic composition as a key determinant of nanoparticle toxicity, demonstrating that small changes in LDH composition can substantially alter their biological effects. From an applied perspective, these results emphasize the importance of considering environmental safety alongside delivery performance in the design of nanocarriers for agricultural applications. Therefore, this study contributes to the development of safer LDH-based RNAi delivery systems and reinforces the need for multispecies ecotoxicological assessments when evaluating nanomaterials intended for agricultural use.

## 6. Author Contributions

A. L. F., T. C. M. S., C. C. C., and G. C. C. contributed to conceptualization, methodology, investigation, writing the original draft, and review & editing. M. A. M., H. C. O., R. V. R., and M. B. J. contributed to review & editing, supervision, and funding acquisition. All authors read and approved of the final manuscript.

## 8. Data availability

All data supporting this study are included within the article and its figures and tables.

## 9. Acknowledgements

This study was funded by the Coordination for the Improvement of Higher Education Personnel – Brazil (CAPES) – Funding Code 001; the São Paulo Research Foundation (FAPESP; grants no. 2023/05204-5, 2023/06143-0, 2022/03250-7, 2024/18680-2, 2021/08354-2, 2022/16724-7, and 2020/01218-3); the National Council for Scientific and Technological Development (CNPq); and the Support Fund for Teaching, Research, and Extension (FAEPEX-UNICAMP; grants no. 2633/17, 2421/20, 2237/21, 2178/22, and 2534/23). RVR is a CNPq fellow (Grant no. 304295/2022-1).

