## Supplemental material for "Organism-dependent ecotoxicological effects of transition metal-containing layered double hydroxides"

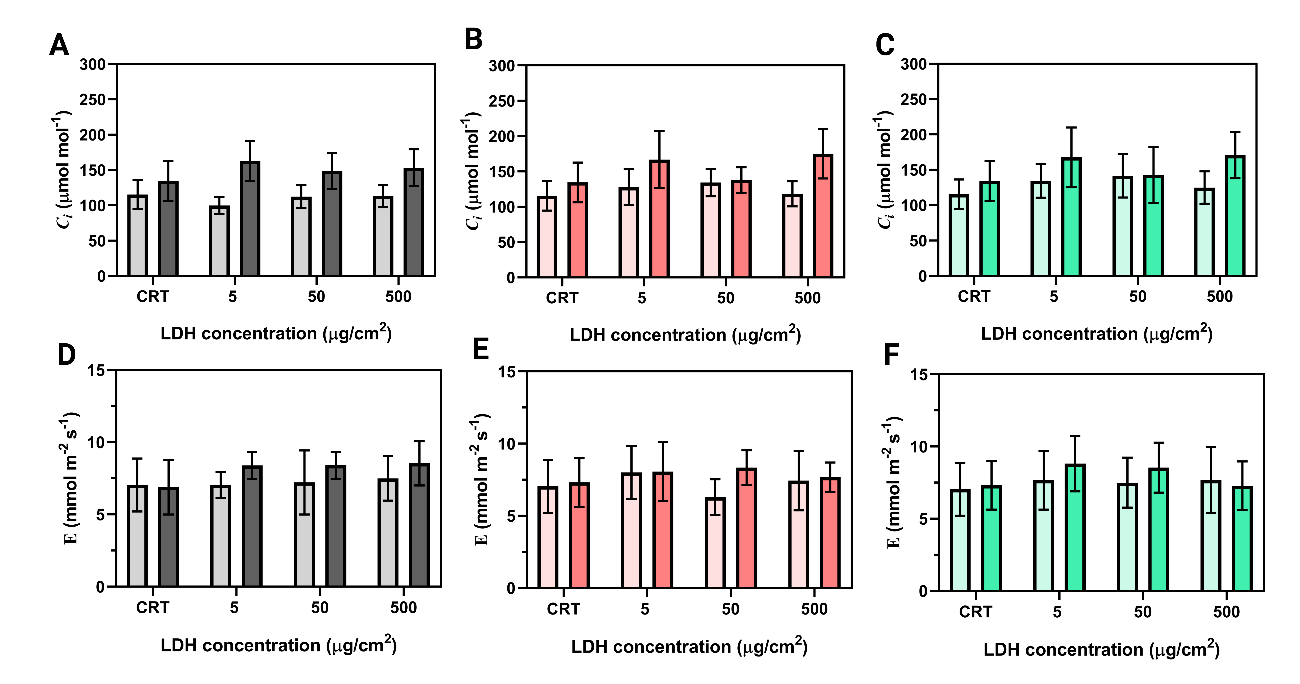

**S1: Physiological responses of maize plants exposed to layered double hydroxide (LDH) nanoparticles with different metal compositions (Al-Ni, Al-Co, and Al-Cu).** (A–C) Internal $\text{CO}_{2}$ concentration ($C_{i}$), and (D–F) transpiration rate ($E$) measured in plants treated with Al-Ni, Al-Co, and Al-Cu LDHs, respectively. Data are presented as mean $\pm$ standard deviation (n=7 plants per treatment).

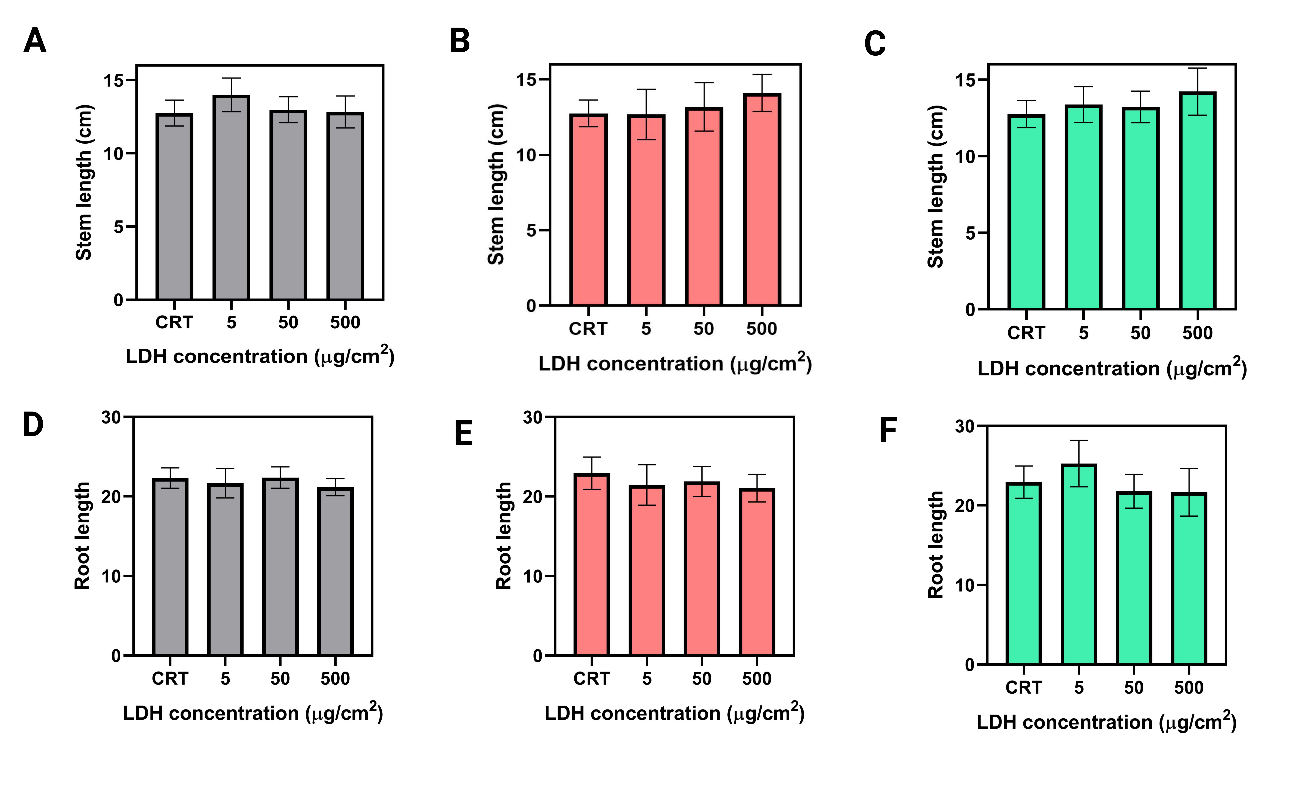

S2: **Biometric responses of maize plants exposed to layered double hydroxide (LDH) nanoparticles with different metal compositions (Al-Ni, Al-Co, and Al-Cu).** (A–C) Stem length, and (D–F) root length measured in plants treated with Al-Ni, Al-Co, and Al-Cu LDHs, respectively. Data are presented as mean $\pm$ standard deviation ($n=7$ plants per treatment).

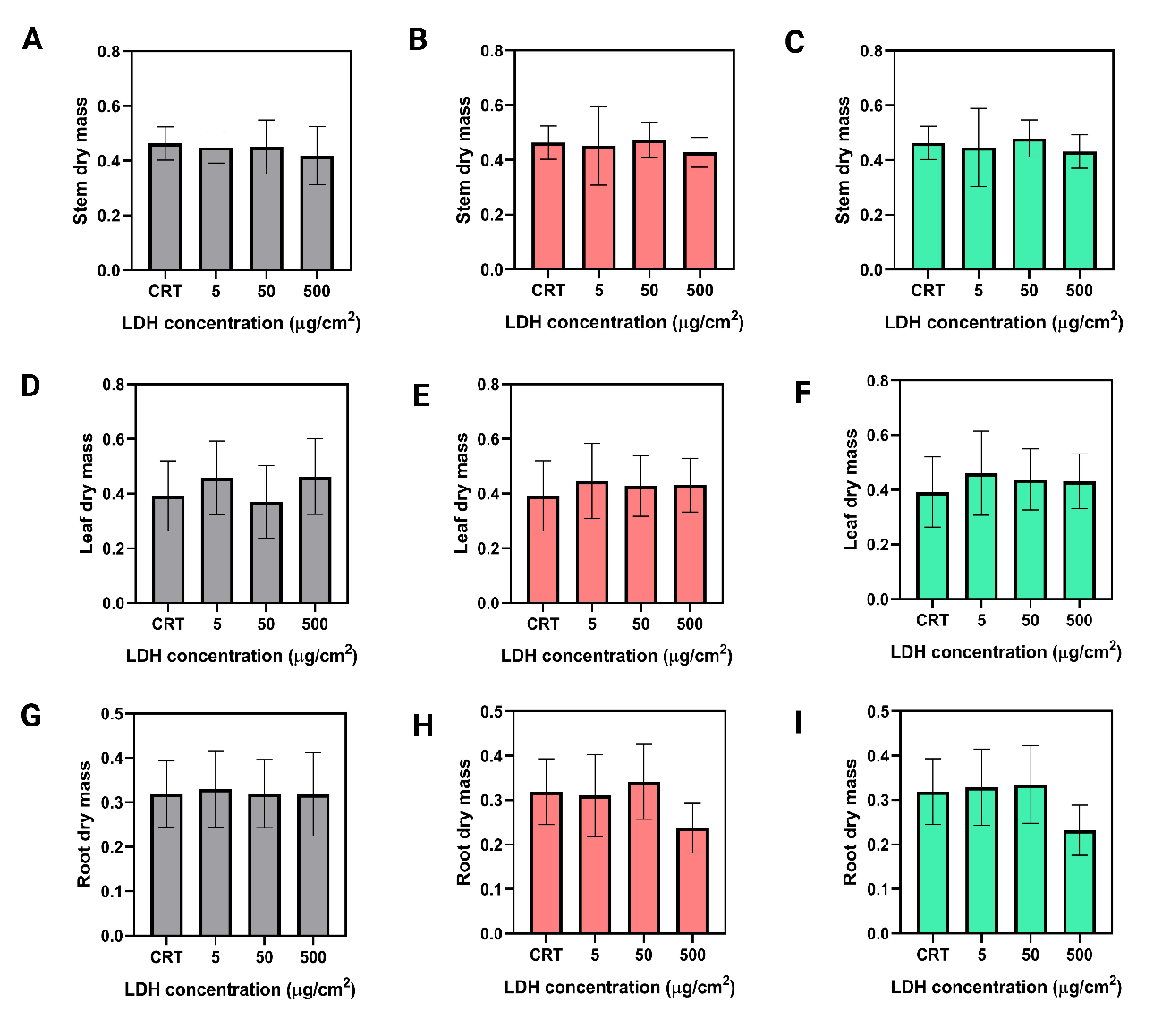
**S3:** **Biometric responses of maize plants exposed to layered double hydroxide (LDH) nanoparticles with different metal compositions (Al-Ni, Al-Co, and Al-Cu).** (A–C) Stem dry mass, (D–F) leaf dry mass and (D-H) root dry mass measured in plants treated with Al-Ni, Al-Co, and Al-Cu LDHs, respectively. Data are presented as mean $\pm$ standard deviation (n=7 plants per treatment).

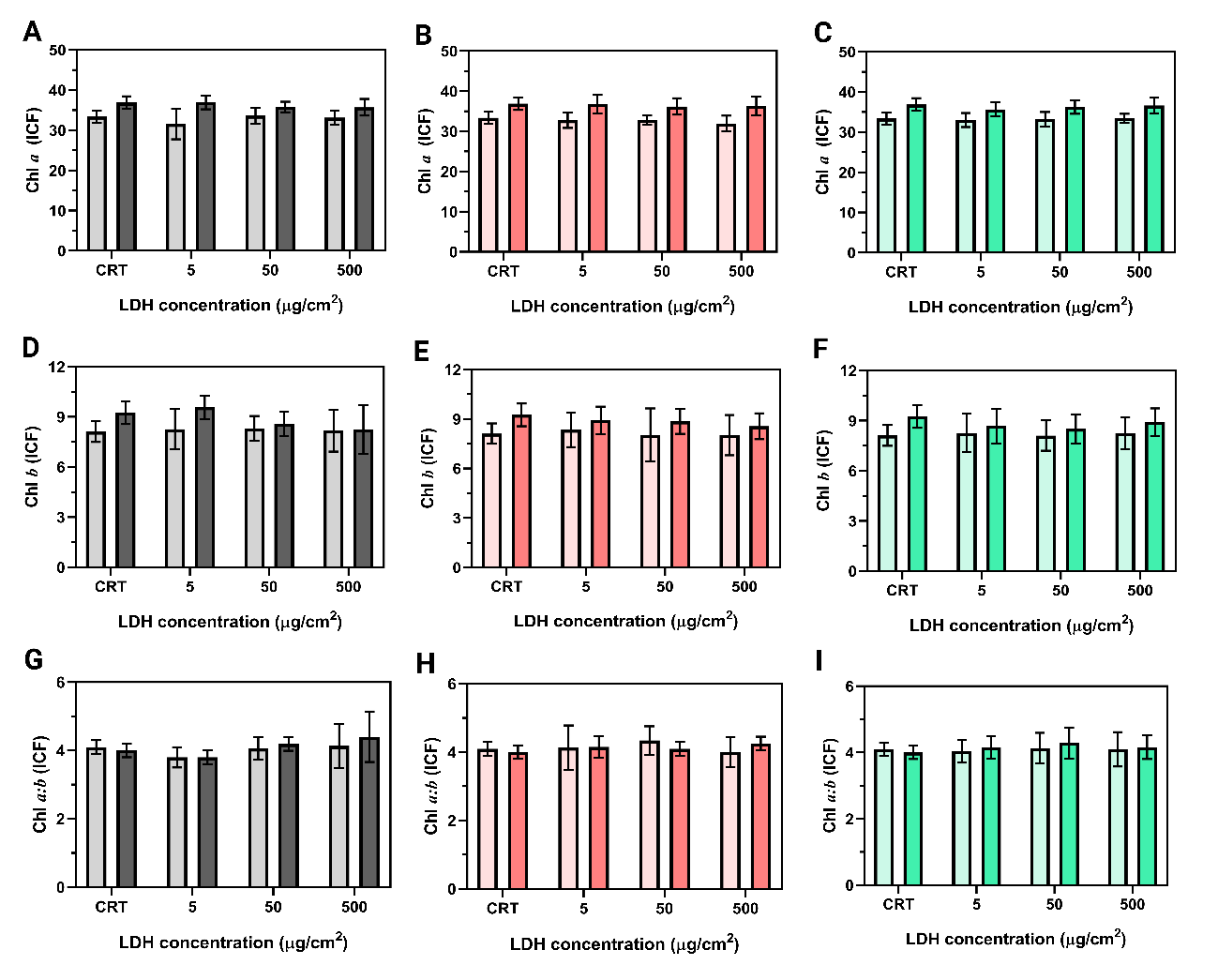

**S4:** **Physiological responses of tomato plants exposed to layered double hydroxide (LDH) nanoparticles with different metal compositions (Al-Ni, Al-Co, and Al-Cu).** (A–C) Total chlorophyll *a*, (D–F) total chlorophyll *b*, and (G–I) chlorophyll *a:b* ratio measured in plants treated with Al-Ni, Al-Co, and Al-Cu LDHs, respectively. Data are presented as mean ± standard deviation (n = 10 plants per treatment).
